# Reconciling age-related changes in behavioural and neural indices of human perceptual decision making

**DOI:** 10.1101/228965

**Authors:** David P. McGovern, Aoife Hayes, Simon P. Kelly, Redmond O’Connell

## Abstract

Ageing impacts on decision making behaviour across a wide range of cognitive tasks and scenarios. Computational modeling has proven highly valuable in providing mechanistic interpretations of these age-related differences; however, the extent to which model parameter differences accurately reflect changes to the underlying neural computations has yet to be tested. Here, we measured neural signatures of decision formation as younger and older participants performed motion discrimination and contrast-change detection tasks, and compared the dynamics of these signals to key parameter estimates from fits of a prominent accumulation-to-bound model (drift diffusion) to behavioural data. Our results indicate marked discrepancies between the age-related effects observed in the model output and the neural data. Most notably, while the model predicted a higher decision boundary in older age for both tasks, the neural data indicated no such differences. To reconcile the model and neural findings, we used our neurophysiological observations as a guide to constrain and adapt the model parameters. In addition to providing better fits to behaviour on both tasks, the resultant neurally-informed models furnished novel predictions regarding other features of the neural data which were empirically validated. These included a slower mean rate of evidence accumulation amongst older adults during motion discrimination and a beneficial reduction in between-trial variability in accumulation rates on the contrast-change detection task, which was linked to more consistent attentional engagement. Our findings serve to highlight how combining human brain signal measurements with computational modelling can yield unique insights into group differences in neural mechanisms for decision making.

---

Ageing is associated with a progressive decline in many aspects of cognitive function including episodic memory^1^, working memory^2^, speed of processing^3^ and task-switching^4^. In an effort to better understand these wide-ranging changes, researchers have sought to pinpoint core mental operations that are impacted by ageing (e.g. refs.^3,5,6^). One such operation is perceptual decision making, the process whereby sensory information is translated into goal-directed actions.

Much of what we know about how ageing affects perceptual decision making comes from behavioural studies employing sequential sampling models,^7–9^. The core principle underlying these models is that noisy sensory evidence is accumulated until a predetermined quantity has accrued in favour of one of the decision alternatives. The two most consistent findings to emerge from this work have been an age-dependent widening of decision boundaries^10–16^, consistent with older adults adopting more cautious decision policies^17^ and an age-related slowing of non-decision processing indicating a delay in sensory encoding and/or motor execution^10–12, 14, 18–21^. Perhaps surprisingly, ageing effects on the parameter that dictates the mean rate of evidence accumulation (‘drift rate’) vary substantially across studies and appear to be highly task-dependent (e.g. refs.^10,14,21,22^). For instance, whereas older and younger adults had similar drift rates on a signal detection task^10^, older adults had lower drift rates on a letter discrimination task^21^ and higher drift rates during motion discrimination^16^.

While these mathematical models have provided valuable insights into the effects of ageing on decision making, there are limits on what can be gleaned from the modelling of behavioural data alone (see ^23^ for a recent review). For instance, it is difficult to ascertain whether any age-related differences in drift rate emanate from differences in the quality of sensory encoding, the integrity of the evidence accumulation process itself or engagement of other systems that play a supporting role in perceptual decision making, such as neuromodulatory and attention systems which are known to be affected by ageing (e.g. ref. ^24^). More generally, abstract decision models that quantitatively capture behaviour do not necessarily reflect the neural computations underlying decision formation, as evidenced in recent non-human primate research ^25–27^. Some animal neurophysiology studies have thus begun to use neural signatures of decision formation directly to construct novel model variants that are more representative of the neural implementation of the decision process as well as its behavioural output^25,28^. The recent identification of analogous decision signals in non-invasive recordings presents an opportunity to similarly develop neurally-informed models of human decision making. These human brain signals bear all of the same decision-predictive characteristics as have been reported for build-to-threshold decision signals observed in single-unit recordings^29–31^, including a gradual buildup whose rate predicts reaction time and is proportional to evidence strength, and a fixed amplitude immediately prior to decision reports consistent with a boundary crossing effect^32–34^. Two functionally-distinct categories of human decision signals have been characterised: effector-selective signals that represent the translation of sensory evidence into a specific motor plan, such as lateralised oscillatory activity in the beta-band^33–36^, and a domain-general signal found in the event-related potential, termed the centroparietal positivity (CPP), that exhibits the same evidence accumulation properties irrespective of whether a response is immediate, delayed^37^ or not required at all^33^. The buildup of the CPP reliably precedes evidence-selective motor preparation signals^32^, suggesting that it reflects a processing level that intermediates between sensory encoding and motor preparation.

To investigate the impact of ageing on the neural decision process and to examine correspondences with the predictions derived from behavioural modelling, we asked a group of younger and older participants to perform a continuous version of a random dot motion task^32^ and a gradual contrast-change detection task^33^, while we recorded 64-channel electroencephalography (EEG). This approach allowed us to isolate both effector-selective and domain-general indices of decision formation whose dynamics we compared to key parameter values derived from fitting the most popular sequential sampling variant, the drift diffusion model^9^, to the behavioural data. For the continuous random dot motion task, the diffusion model accounted for age-related performance deficits (longer reaction times and more misses) in terms of a widening of decision boundaries in older adults. However, there were no differences in the amplitude of either the effector-selective or domain-general decision signals at the time of the decision report. Instead, older adults showed slower decision signal buildup of neural evidence accumulation signals, which is equally consistent with the performance deficits in this task. Meanwhile, in the contrast-change detection task, older subjects performed better (less variable reaction times and fewer misses), and while the model fits explained this by an increase in both drift rate and decision bound, no age-related differences in the corresponding neural signal measurements were observed. We go on to show that if the diffusion model is constrained to take account of these neurophysiological observations, the model provides a better fit to the motion discrimination data and a comparable fit to behaviour on the contrast-change detection task despite having fewer free parameters, as well as producing estimates of the non-constrained parameters that accord with the corresponding neural measures. Specifically, when decision bounds were constrained to be equal and starting-point variability was added as a free parameter to take account of premature evidence accumulation observed in older adults on the motion task, the model was then able to capture the drift rate difference evident in the neural data. In the contrast-change task, reduced across-trial variability in drift rate in older adults that became apparent in the neurally-constrained model was reflected in reduced variability both in the buildup rate of the decision signals and in pretarget alpha-band activity, suggesting more consistent attentional engagement. Together, our data suggest that human neurophysiological signals can play an important role in constraining models of perceptual decision making and revealing key mechanistic differences that may go undetected using behavioural modelling alone.

## Results

### Motion Discrimination

#### Behaviour

In a first experiment, 35 younger (17 males, age range: 18-38, mean age: 22 years old) and 31 older participants (13 males, age range: 66-83, mean age: 74 years old) performed a continuous version of a random dot motion task in which they were required to discriminate the direction of intermittent periods of coherent motion (randomly set to 30 or 60%) that occurred within a continuous stream of incoherent motion (Figure 1a; see also ref. ^32^). For both young and old participants, higher sensory evidence strength (motion coherence) resulted in fewer misses (p<0.001 for both age groups, Wilcoxon signed-rank test) and faster reaction times (RTs; p<0.001 for both age groups, Wilcoxon signed-rank test). In keeping with previous studies examining the effects of ageing on motion discrimination^16,38,39^, older adults were significantly less accurate than their younger counterparts for both levels of motion coherence (Figure 1b; 30%: U=250, p<0.001, 60%: U=167, p<0.001, Mann-Whitney test) and also displayed significantly slower RTs (Figure 1b; 30%: U=263, p<0.001, 60%: U=114, p<0.001, Mann-Whitney test). Further analysis revealed that older adults made more erroneous responses (30% coherence: 2.6% ± 5.4 vs. 0.5 ± 1.2%, U=357, p=0.005; 60%: 5.4% ± 6.6 vs. 0.8% ± 1.1, U=199, p<0.001) and more misses (30%: 6.6% ± 10.3 vs. 1.3 % ± 2.1, U=344, p=0.007; 60%: 0.4% ± 1.1 vs. 0% of targets, U=473, p=0.04) than their younger counterparts, while there were no differences in the false alarm rates between the two age groups (Younger (mean ± SD): 0.39 ± 0.5 per block, Older: 0.54 ± 0.7 per block, U=497, p=0.55, Mann-Whitney test).

**Figure 1:**
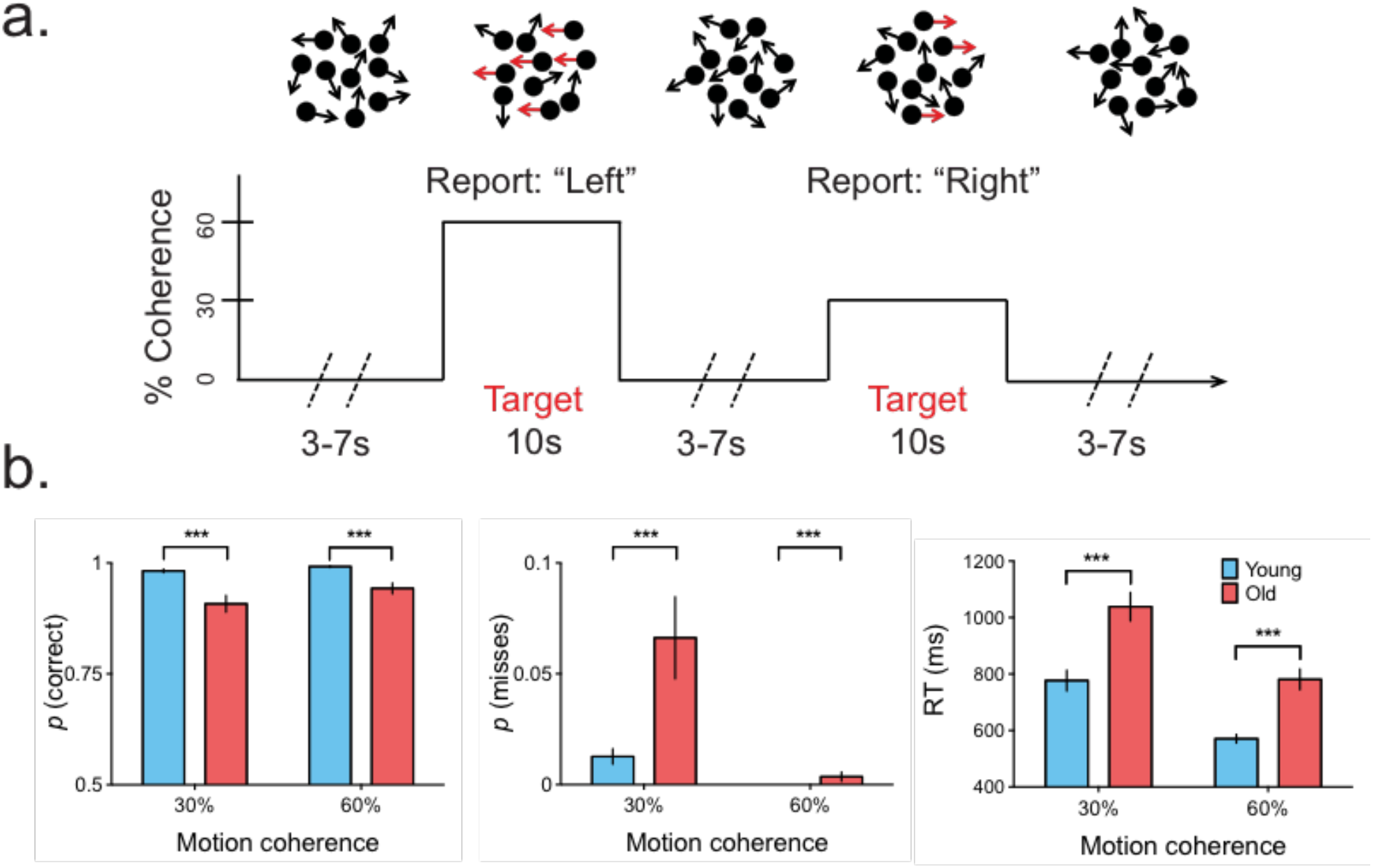
Random dot motion task and associated behaviour. a) Task schematic of the continuous random dot motion task. Participants monitored a centrally-presented dot pattern for transitions from random to coherent motion. b) Older adults were slower and made more incorrect responses at discriminating motion direction than their younger counterparts. Error bars indicate ±1 SEM.

To gain a better understanding of what aspects of the decision process led to these differences in behaviour, we fit a drift diffusion model to the accuracy and RT data for both age groups (Figure 2a). We first identified the key parameters required to fit the model to data pooled across participants from each group (decision bound, drift rate, non-decision time, across-trial drift rate variability; see Supplementary Fig. 1 and *Methods* for further information on our approach), before fitting the model to each individual’s data from the two groups (Figure 2c). An analysis of incorrect responses concluded that they were highly unlikely to have arisen from incorrect decision boundary crossings through evidence accumulation (see *Methods* for further details) and therefore incorrect responses were omitted from the fitting procedure.

**Figure 2:**
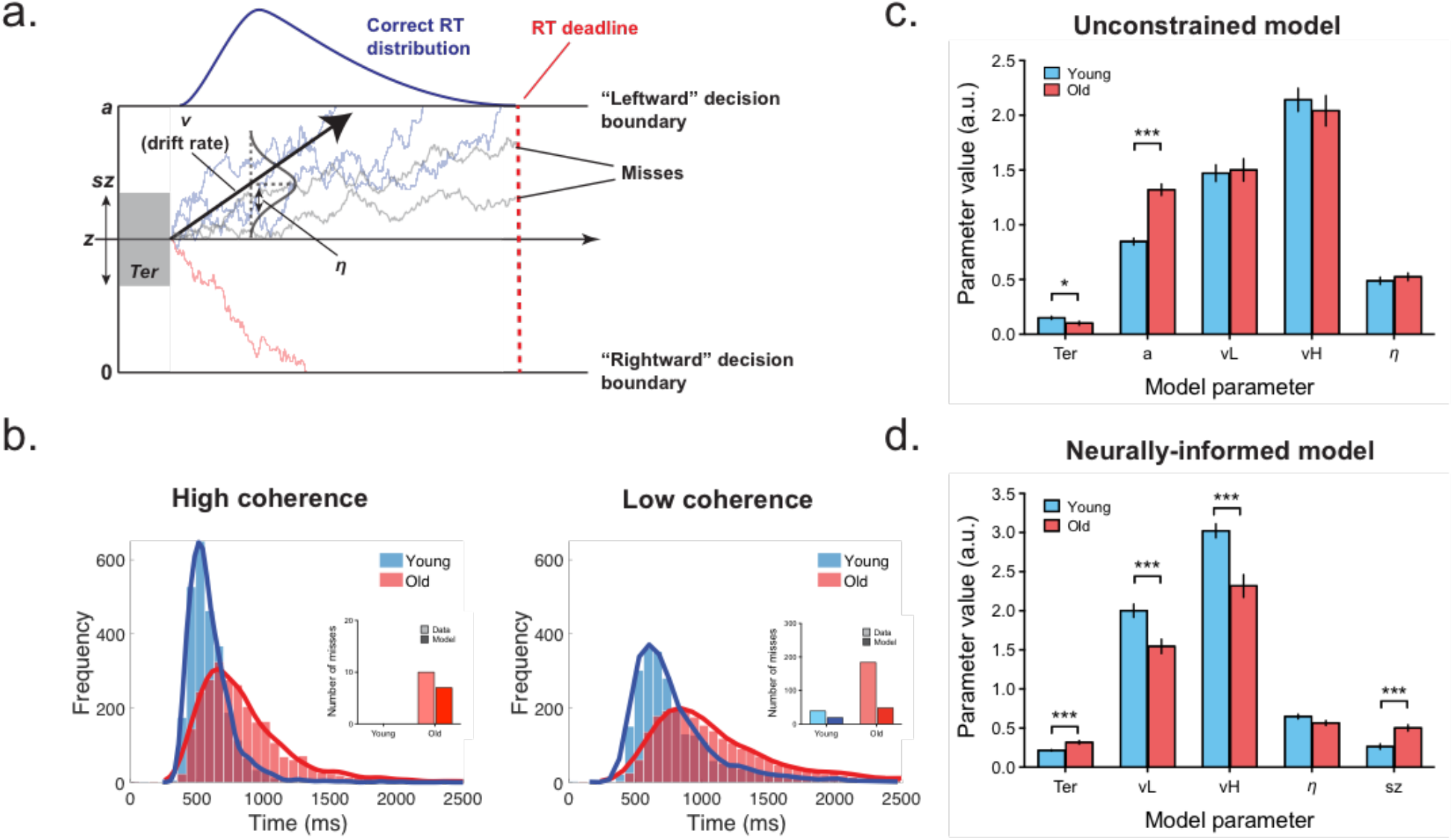
Comparison of diffusion model fits to motion discrimination data with decision boundary free to vary and constrained. a) Schematic of the two-choice drift diffusion model that was fit to the data. Evidence accumulation begins at starting point, *z*, and a response is initiated when the cumulative evidence reaches the correct, *a*, or incorrect, 0, response boundary. The mean rate of accumulation is determined by the drift rate, *v*, which can vary from trial-to-trial. This variability is assumed to be normally distributed with a standard deviation of *η*. In the neurally-informed model, the starting point of the accumulation process could also vary on a trial-to-trial basis, *sz*, and this variability was assumed to be uniformly distributed. On some trials, the cumulative evidence does not reach either decision boundary before the response deadline and in these cases the trial is classified as a miss. b) Neurally-informed model fits (solid lines) to observed reaction time distributions (histograms) pooled across participants. Inset of each panel: Observed and model estimates of target misses. c) Mean parameter estimates derived from model fits to individual data where decision boundary was free to vary. d) Mean parameter estimates derived from model fits to individual data where decision boundary was constrained to be the same across age groups and where starting point variability was added as a free parameter.

The model produced good fits to the pooled group data for both the younger (G^2^ = 51.51) and older (G^2^ = 70.81) groups. Consistent with previous behavioural modelling studies of motion discrimination and other two-alternative perceptual decisions^10,16,21^, both the group-level and individual fits suggested that older adults had significantly larger boundary separations than younger adults (Figure 2c; U=91, p<0.001, Mann-Whitney test; see also Supplementary Fig. 1b), with no significant differences in drift rate (Figure 2c; *vL:* t(64)=0.23,p=0.82; *vH*: t(64)=0.58, p=0.56). Older adults also had significantly shorter non-decision times (t(64)=2.1, p=0.04).

#### Neurophysiology

To measure the impact of ageing on neural evidence accumulation processes, we recorded EEG while participants performed the random dot motion task. In line with our earlier findings^32,40^, the onset of coherent motion elicited a rising positivity in the event-related potential over centroparietal scalp regions in both younger and older participants (Figure 3a). Our previous work has demonstrated that this centroparietal positivity (CPP) exhibits the key properties of bounded accumulation that are central to sequential sampling models of perceptual decision making ^32,33^, while other work has identified similar properties in lateralised oscillatory activity in Mu/Beta band (e.g. ^33–35^). Specifically, these signals exhibit a gradual buildup whose rate scales with evidence strength and reaches a stereotyped amplitude at the time of response. Importantly, both of these signals have also been shown to be sensitive to experimental manipulations previously shown to modulate participants’ decision boundaries, such as predictive cueing paradigms and experiments that manipulate the speed/accuracy emphasis of a task (e.g. ^34,36,41^). Here, we took advantage of these evidence accumulation properties to examine the effect of ageing on the buildup and amplitude of response-locked CPP and oscillatory Mu/Beta activity and compare the findings to our modelling results.

**Figure 3:**
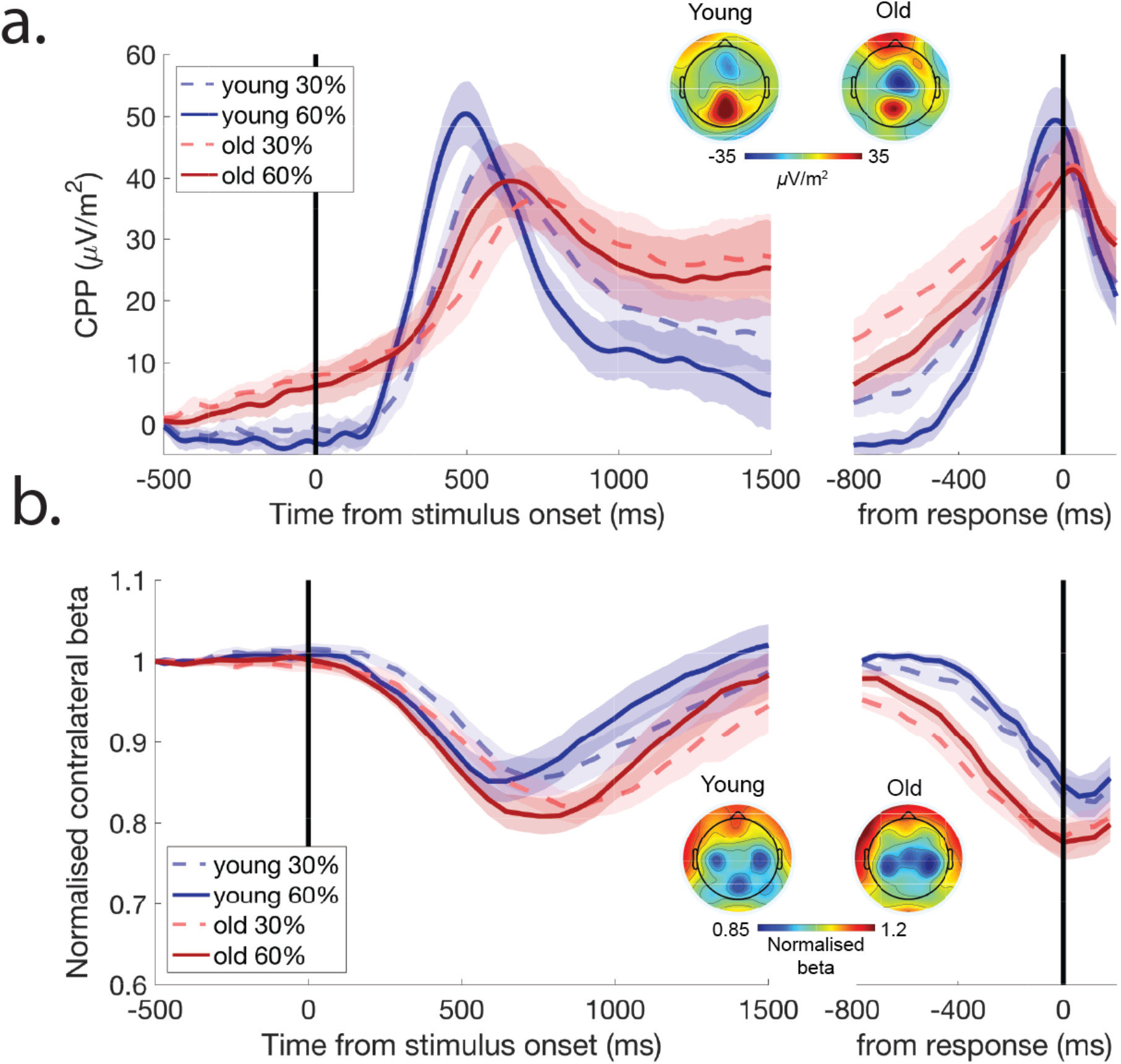
During motion discrimination older adults exhibit a slower rate of evidence accumulation relative to younger adults, with both age groups displaying similar amplitudes at response. a) Stimulus-locked (left) and response-locked (right) CPP waveforms for young and older participants performing the continuous random dot motion task. The rate of CPP buildup differed between the age groups, with a slower rate of accumulation in the older group compared to the young group, while there was no difference in CPP amplitude between the different age groups at the time of decision report (black solid line on right panel). b) A similar pattern of results was observed for contralateral Mu/Beta activity. Shaded areas indicate ±1 SEM of data points.

Initial inspection of the data revealed that the buildup of the CPP of the older group actually commenced before the onset of coherent motion suggesting a tendency towards premature evidence accumulation (see Supplementary Fig. 2). To account for this, we baseline-corrected the ERP waveforms of both age groups relative to a 100 ms interval preceding this initial build-up (600 to 500 ms prior to coherent motion; see *Discussion* for further consideration of this analysis step).

Across both groups, the CPP shared two of the key characteristics of the theoretical decision variable central to sequential sampling models, namely a gradual buildup whose rate increased in proportion to evidence strength (i.e. motion coherence, F(1,64)=25.13, p<0.001, BF_01_<0.001), and a stereotyped amplitude immediately prior to response execution that did not differ as a function of coherence (F(1, 64)= 0.8078, p=0.37, BF_01_=3.53). The rate of CPP buildup differed between the age groups, with a slower rate of accumulation in the older group compared to the young group (F(1, 64) = 10.73, p=0.002, BF_01_=0.005), while there was no significant difference between age groups in CPP amplitude at response (F(1, 64) = 1.72, p=0.19). Although our Bayes factor analysis indicated that the evidence was inconclusive as to whether there was a significant difference in the CPP amplitude at response (BF01=1.42), it should be noted that the numerical trend favoured a *lower* CPP amplitude in the older group, and is thus at odds with the prevailing view to emerge from the modelling literature that older adults implement higher decision bounds (e.g. ^10–16^). A qualitatively similar pattern of results was observed at the level of effector-selective motor preparation, with contralateral Mu/Beta activity (Figure 3b) reaching a fixed amplitude before response execution irrespective of evidence strength (F(1, 64)= 0.61, p=0.44, BF_01_=3.9), although it was again inconclusive as to whether the amplitude at response varied as function of age group with anecdotal evidence in favour of there being no difference (F(1,64)=2.76, p=0.1, BF_01_=1.84). Also in keeping with the CPP results, the buildup rate of contralateral Mu/Beta was faster for higher motion coherence (F(1,64)=10.85, p=0.002, BF_01_=0.007) and in younger adults (F(1,64)=5.9, p=0.018, BF01=0.34).

#### Neurally-informed modelling

Our neurophysiological data indicate that age-related decrements in motion discrimination performance arise due to slower accumulation of motion information in older adults, with little or no difference in the decision boundary positions of younger and older participants. However, these observations are at odds with our behavioural modelling results and those of previous studies (e.g. ref. ^16^). In an effort to address this discrepancy, we fit an additional model to the behavioral data in which, based on our neurophysiological observations, the decision bound was fixed to the mean of the young and older group parameter values from the original group-level model fits (see *Methods*). To account for the early buildup of the CPP observed in older adults (see Figure 3a), we also included starting point variability as a free parameter in the model on the grounds that it would capture any variance in performance and decision-signal buildup due to premature evidence accumulation (e.g. ref. ^42^) irrespective of the particular accumulation strategy that was adopted (e.g. leaky versus non-leaky). The model fits from this neurally-informed model were 22 better than the original model for both the younger (G^2^ = 46.9) and older (G^2^ = 54.3) groups despite both models having the same number of free parameters. Moreover, the age group differences in the parameter values from the neurally-informed model were more in keeping with our neurophysiological data, with significantly lower drift rates in older subjects for both low and high coherence stimuli (Figure 2d; *vL*: U=264, p<0.001, *vH*: U=242, p<0.001, Mann-Whitney tests; see also Supplementary Fig. 1c). Older adults also displayed significantly higher levels of starting point variability compared to their younger counterparts (t(64)=4.49, p<0.001), which accords with the observation of larger pretarget CPP build-up in this group; since the CPP manifests as a positive deflection irrespective of which decision alternative the evidence favours, its average amplitude at target onset should be larger when there is greater starting point variability. Between-trial drift rate variability was also higher in younger participants than older participants, although this effect was not statistically significant (t(64)=1.94, p=0.056; see *Contrast-change detection task* for further analysis and discussion). We also note that even when starting point variability was not free to vary between groups, a model with constrained bounds produced more parsimonious fits (BICs=93.29 and 108.37 for younger and older groups, respectively) than a model with unconstrained bounds (BICs=95.17 and 113.69) and again highlighted significantly higher drift rates in older participants (*vL*: U=220, p<0.001, *vH*: U=182, p=0.001, Mann-Whitney tests). Thus, adapting the diffusion model according to our neurophysiological observations produced better fits to behaviour while also providing a better account of the age-related differences observed in our neural measures of evidence accumulation.

### Contrast-change Detection

#### Behaviour

To assess whether the age-related effects we observed on the random dot motion task generalised to other perceptual decision making tasks, the same participants performed a contrast-change detection task^33^ within the same testing session, with the order of the tasks counterbalanced across participants. Participants were required to continuously monitor a checkerboard stimulus for intermittent reductions in contrast, which were reported via a right-hand button click (Figure 4a). Older adults surprisingly outperformed their younger counterparts on this task, detecting more of the contrast reductions (Figure 4b; U=456, p=0.005, Mann-Whitney test) despite maintaining similar average response times (Figure 4b; t(74)=0.179, p=0.86) and false alarm rates (Younger: 1.6 per block ± 1.4, Older: 1.6 per block ± 1.9, U=636, p=0.37, Mann-Whitney Test). The younger group did, however, display increased RT variability compared to the older group (Figure 4c; Mean coefficient of variation across participants: Younger= 0.234, Older=0.209, t(74)=2.5, p=0.015). To establish what aspects of the decision process gave rise to the older adults outperforming their younger counterparts on this task, we fit the data with a one-choice drift diffusion model (Figure 5a; ref. ^43^), with the same free parameters as in the analysis of the motion discrimination (decision bound, drift rate, non-decision time, between-trial drift variability). A key difference in this model, however, was that we assumed that the drift rate rose linearly over the duration of the target stimulus to reflect the ramping evidence signal (i.e. decreasing contrast; see *Methods*). This model provided a very good fit to the pooled group data (Supplementary Fig. 3a) for both the younger (G^2^ =84.24) and older (G^2^ =87.33) groups. The parameter estimates from the fits of this model to the individual data indicated that the behavioural advantage in the older group was linked to the combined effect of two distinct parameter differences: higher drift rates (Figure 5c; t(74)=3.26, p=0.002; see also Supplementary Fig. 3b), as well as wider decision boundary separations than young adults (t(74)=2.4, p=0.02).

**Figure 4:**
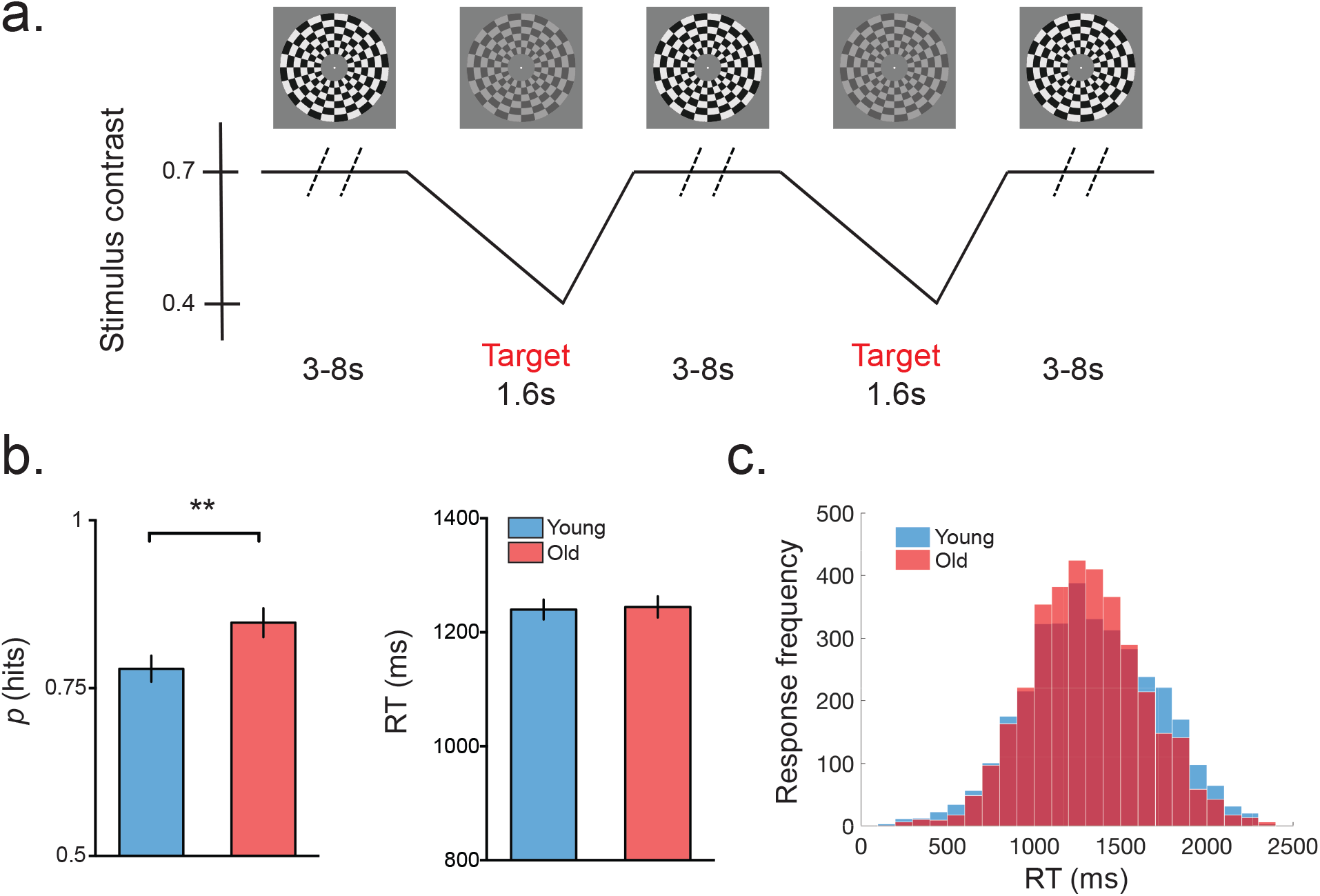
Gradual contrast-change detection task and associated behaviour. a) Schematic of contrast-change detection task. Participants monitored a checkerboard annulus for intermittent decreases in stimulus contrast. b) Older adults detected more of the contrast targets than younger adults, while maintaining similar reaction times. c) Reaction time distributions for young and older age groups.

**Figure 5:**
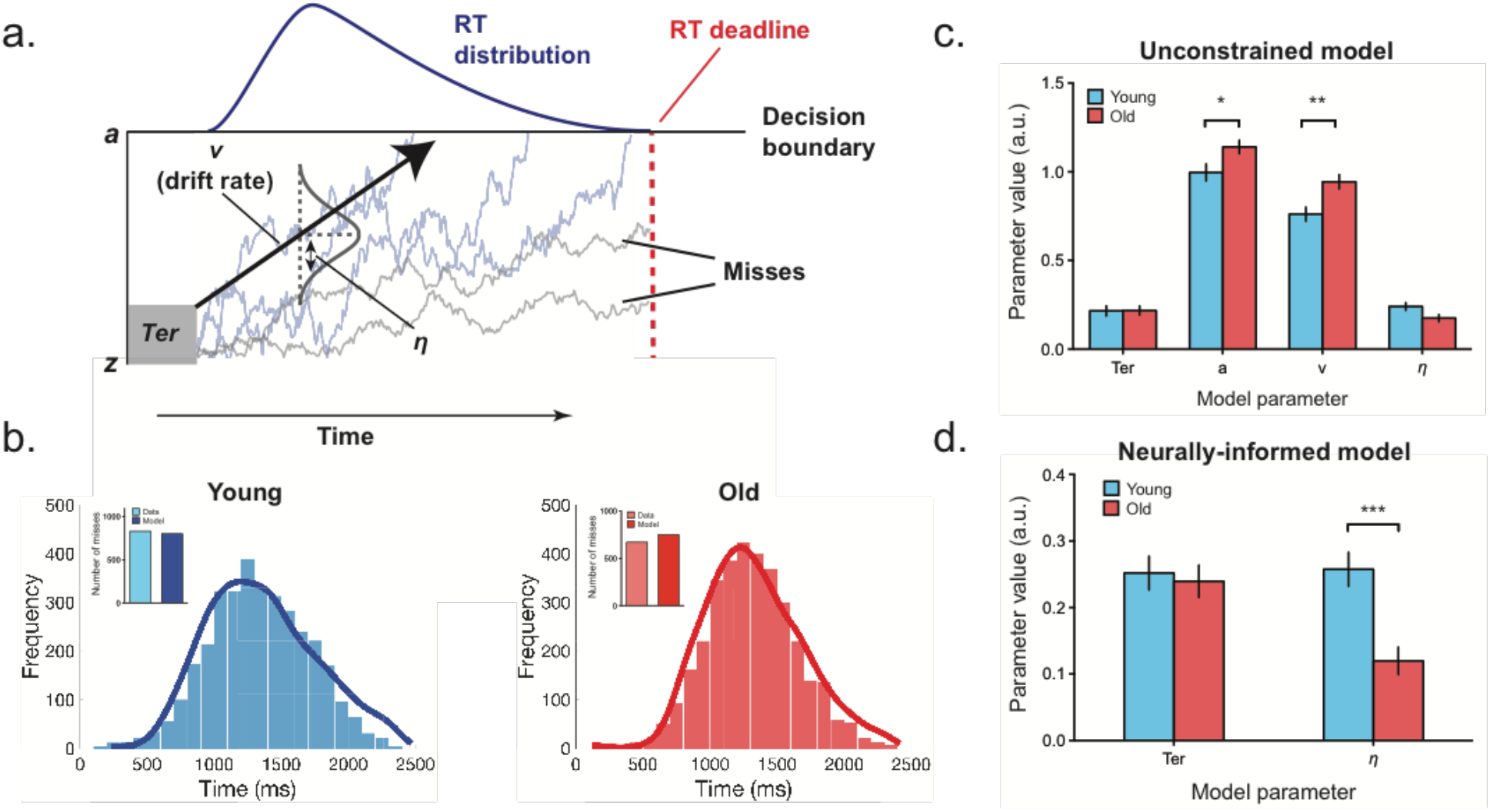
Comparison of diffusion model fits to contrast-change detection data with decision boundary and drift rate free to vary and constrained. a) Schematic of the one-choice drift diffusion model that was fit to the data. Evidence accumulation begins at starting point, *z*, and a response is initiated when the cumulative evidence reaches the decision boundary, *a*. The rate of accumulation is called the drift rate, *v*, which can vary from trial-to-trial. This variability is assumed to be normally distributed with a standard deviation of *η*. On some trials the cumulative evidence does not reach the decision boundary before the response deadline and in these cases the trial is classified as a miss. b) Neurally-informed model fits (solid lines) to observed reaction time distributions (histograms) pooled across participants. Inset of each panel: Observed and model estimates of target misses. c) Mean parameter estimates derived from model fits to individual data where decision boundary and drift rate were free to vary. d) Mean parameter estimates derived from model fits to individual data where decision boundary and drift rate were constrained.

#### Neurophysiology

To examine whether comparable effects were observed in the neural indices of decision formation, we again measured the CPP and left hemisphere beta (LHB) as assays of sensory evidence accumulation and motor preparation, respectively. In addition to these measures, the gradual contrast detection task also allowed us to track the basic sensory encoding of contrast over time by measuring the amplitude of the occipital steady-state visual evoked potential (SSVEP; refs. ^33,44^) generated by the on-off flicker of the checkerboard stimulus. Although the amplitude of the SSVEP was greatly reduced in the older group compared to the young group, when the signals were normalised to take account of these baseline differences (see *Methods* for further details), the SSVEPs for both age groups reliably tracked the decrease in stimulus contrast displaying remarkably similar stimulus- and response-aligned waveforms (Figure 6a). Furthermore, both the CPP (Figure 6b) and LHB (Figure 6c) were very similar across age groups, with no significant differences in the buildup rates (CPP: t(74)=1.18, p=0.24, BF_01_=3.02; LHB: t(74)=0.99, p=0.32, BF_01_=3.64) or peak amplitudes (CPP: t(74)=0.68, p=0.5, BF_01_=4.62; LHB: t(74)=0.16, p=0.88, BF_01_=5.66) of these signals in the response-aligned waveforms.

**Figure 6:**
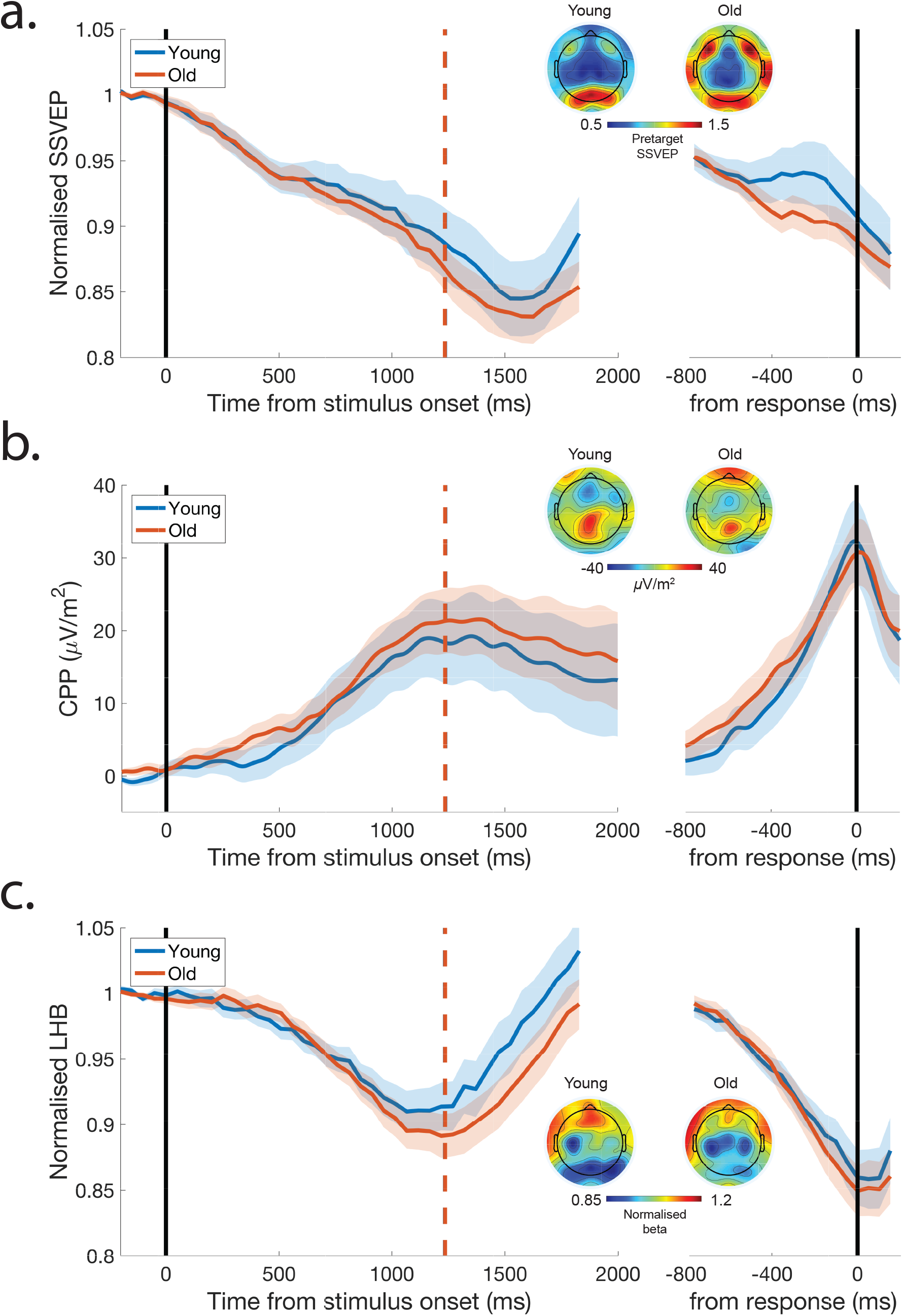
Sensory encoding, decision formation and motor preparation signals exhibit similar dynamics in young and older participants on the contrast-change detection task. a) Stimulus-locked (left) and response-locked (right) SSVEP displays similar buildup rates and amplitudes at response for both age groups. A similar pattern was observed for both the CPP (b) and for left hemisphere beta (LHB) band activity (c). Dashed lines on stimulus-locked CPP and LHB indicate mean reaction times of both age groups.

#### Neurally-informed modelling

Our initial neural signals analysis did not uncover any differences at the sensory encoding, decision formation or motor preparation processing levels that could account for the enhanced contrast detection performance of the older participants. As in the random dot motion data, these electrophysiological data appear to be at odds with our findings from a fit of a standard model, which suggested that older adults displayed higher rates of evidence accumulation and elevated decision boundaries. In an attempt to reconcile these findings, we fixed the boundary criterion and drift rate parameters to intermediate values between the younger and older values derived from the initial model fit (4 free parameters) and fit the resulting model (2 free parameters) to the behavioural data (Figure 5b). While the fits of this reduced model to the pooled data were marginally worse compared to the unconstrained model fits for both younger (G^2^=97.31) and older (G^2^=90.72) groups, the BIC values indicated that the constrained model (Younger: 113.79, Older: 107.2) provided a more parsimonious fit to the data compared to the original model (Younger: 117.19, Older: 120.3) with 4 free parameters. Fitting this neurally-constrained model to the individual data suggested that the key factor underlying the age-related behavioural advantage was a higher degree of between-trial drift rate variability in younger adults compared to older adults (Figure 5d; U=359, p<0.001, Mann-Whitney test; see Supplementary Fig. 3c for parameter values from fits to pooled data). In the following section we sought evidence in our neurophysiological data to support these model findings.

#### Linking neurally-informed modelling to neurophysiology

The results from the neurally-constrained diffusion model suggest that the main factor driving the older adults’ behavioural advantage on the contrast-change detection task is reduced between-trial drift rate variability. To establish whether there was a corresponding age-related reduction in the variability of the neural signatures of decision formation, we calculated single-trial estimates of the CPP slope for the two age groups. This analysis revealed that the buildup rate of the CPP was significantly more variable in the young group compared to the older group (Mean SD across participants: Young=0.31, Old=0.21, t(74)=5.7, p<0.001) in keeping with the modelling results. Furthermore, a similar age-related reduction in the variability of the buildup rate of the CPP was also observed in the data from the random dot motion task (Median SD across participants: Young=0.16, Old=0.13, U=385, p=0.04, Mann-Whitney Test) suggesting that this effect generalises across different perceptual tasks.

One potential reason for the age-dependent decrease in between-trial variability of evidence accumulation is that older adults maintained a greater level of engagement with the task (e.g. ^16^), while younger participants may have been more susceptible to fluctuations of attention over the course of the experiment. Although the current experiment did not directly manipulate the effect of attention on task performance, the amplitude of pretarget alpha-band activity over posterior scalp sites is a well-known electrophysiological signature of endogenous attention^32,45,46^. In line with the idea that younger adults may be more susceptible to fluctuations of attention, across-trial variability of alpha band activity (coefficient of variation, CV) was higher for younger participants compared to their older counterparts (Young: 0.231, Old: 0.185, t(74)=3.14, p=0.002).

## Discussion

Our results highlight ways in which human brain recordings and abstract computational models can be used to reciprocally inform one another to gain deeper insights into the factors that shape perceptual decision making. For both motion discrimination and contrast-change detection decisions, the initial behavioural modelling and neural signal measurements pointed to very different explanations for age-related changes in decision making performance. However, adapting the models to take account of our neurophysiological observations by fixing certain model parameters (i.e. bound for motion discrimination, bound and drift rate for contrast-change detection) and adding in others (i.e. starting point variability for motion discrimination) produced estimates of the remaining unconstrained parameters that could account for the observed ageing effects on the neural decision signals as well as behaviour. Furthermore, the neurally-constrained model fits uncovered additional, novel effects which were in turn confirmed in follow-up neural signal analysis.

Like many other studies to examine the effects of ageing using sequential sampling models^10–16,19,20^, our initial model fits to the behavioural data suggested that the main age-related adjustment in decision making was a widening of the decision boundaries. However, these modelling results ran contrary to our neural signals analysis, which showed no significant age-related differences in the amplitude of either the domain-general or effector-selective signatures of evidence accumulation at the time of decision report. While a Bayes factor analysis suggested that there was strong evidence to indicate a lack of age group differences in the amplitude of either the CPP or Mu/Beta activity at the time of response for the contrast-change detection task, the results for the motion discrimination task were less conclusive. However, it should be noted that the numerical group trends went in opposite directions for these two decision-related signals, with older adults showing smaller CPPs and greater Mu/Beta activity at response compared to their younger counterparts. Thus, across the two experimental tasks we observed little neurophysiological evidence to support the hypothesis that older adults set higher decision boundaries. Importantly, both of these decision-related signals have previously been shown to be modified in a manner consistent with a change in decision threshold via experimental manipulations commonly used to influence decision policies, such as altering the speed/accuracy emphasis of a task and the presentation of predictive cues^34,36,41^, providing confidence that these signals are sensitive enough to detect group differences in decision boundary. Thus, these neurophysiological observations suggest that, in the current experiments, younger and older adults applied similar decision policies. This is not to say that older adults do not exhibit enhanced response caution in some situations. The requirement to continuously monitor stimuli for temporally unpredictable feature changes renders our tasks qualitatively different to those used in previous work in which trials are discretely presented and the stimulus onset provides a cue for the initiation of evidence accumulation. Here, the unpredictable evidence onsets and instructions to respond only when sure likely encouraged the implementation of conservative decision policies in both groups. It may be that older adults do exhibit relatively higher decision boundaries when placed under greater time pressure but our results highlight that any such differences do not generalise to all scenarios.

A further advantage of this modelling approach is that it generated novel predictions regarding the neural data which would otherwise have been overlooked. Our initial analysis of the sensory encoding and decision signals elicited during the contrast-change detection task provided no clear explanation as to why the older adults performed better than younger adults. While the initial model fits to the behavioural data indicated significant age group differences in both drift rate and boundary separation, we did not observe any differences in the corresponding neural measurements (decision signal build-up rates and amplitudes at response, respectively). When the model was constrained to match these neural observations, it accounted for the poorer performance of younger adults via increased trial-to-trial variability in drift rate. Thus, the neurally-constrained model yielded a new, empirically testable prediction that young participants should exhibit greater trial-to-trial variability in the build-up rate of the CPP and, indeed, we found this to be the case during both the contrast and motion discrimination tasks. The increased variability of drift rates and more variable decision signal build-up of the young group were also accompanied by increased variability in posterior alpha-band activity, an established oscillatory marker of attentional engagement^45–47^. These results suggest that older participants were able to maintain attentional engagement more consistently across the duration of the tasks. It cannot be determined here whether this effect reflects a compensatory strategy on the part of the older group or increased motivation to perform well on the task. Alternatively, given the established links between heightened arousal and greater variability in evidence accumulation^48^, it may be that younger adults were more aroused as they completed the experiment. This heightened arousal state could also help to explain the surprising higher miss rate of the younger group; increased drift variability in the absence of other parameter differences should result in elevated miss rates. Regardless of the precise mechanisms underlying the increased drift rate variability of younger adults, these results serve to highlight how combining neurophysiological data with behavioural modelling make it possible to disentangle positive and negative ageing effects.

Our work builds on an increasing trend to use decision-related signals in the human brain to constrain computational models of perceptual decision making (e.g. ^16,49-53^). For instance, Turner and colleagues^49^ have pioneered a powerful neurally-informed computational approach in which model parameters are estimated by simultaneously fitting a model to behavioural data and brain-wide BOLD activations, leading to better model fits and more accurate predictions about withheld behavioural data. This data-driven approach has proven to be particularly beneficial in overcoming the limited temporal resolution of fMRI to isolate decision-related brain structures. Our approach is distinct from that of Turner et al. to the extent that it focuses on the dynamics of a specific set of neural signals that have been empirically verified to trace decision formation in a model-independent fashion^32,33^. Rather than simultaneously fitting the model to the neural and behavioural data, we first fit a standard diffusion model to our behavioural data and the recovered parameter estimates were used to generate predictions about the neural data. We then examined these decision-related neural signals and, in cases where discrepancies arose between the model results and the signal analyses, the neural data was used to constrain the model. In both experiments the constrained model was found to provide a marginally better fit to the behavioural data than the standard model. More importantly, however, the constrained model provided a much better account of the key group differences observed in our neural signatures of decision formation including the slower rate of evidence accumulation and greater noise accumulation at target onset in older adults during motion discrimination, and greater across-trial variability in evidence accumulation rates in younger subjects on the contrast-change detection task.

The continuous monitoring tasks employed in the current study placed a strong emphasis on the detection of stimulus feature changes. In the version of the random dot motion task implemented here, participants frequently missed targets but incorrect discriminations were rare. Consequently, the behavioural data provided fewer constraints for modelling purposes than the discrete trial forced-choice tasks that are more typically employed in the decision making literature. Nevertheless, there is good evidence to suggest that the diffusion model behaves appropriately with these data. First, both the initial and neurally-constrained models provided excellent fits to the behavioural data from both the motion discrimination and contrast-change detection tasks (see Figures 2 & 5, as well as Supplementary Figures 1 & 3). Second, the recovered parameter estimates from the unconstrained models produced age-related effects that are consistent with those reported in previous studies that employed standard two-alternative forced-choice tasks. Most notably, in keeping with the previous literature^10–16^, our modelling results indicated an age-dependent widening of decision boundaries for both the motion discrimination and contrast-change detection tasks. Thirdly, our neurally-informed models also captured all of the key group differences in evidence accumulation dynamics observed in our neurophysiological data. In this sense, the current study highlights how neural data can provide additional constraints for modelling data from behavioural scenarios that are in other ways not ideal for model fitting.

Whereas older adults displayed smaller drift rate parameter estimates coupled with slower mean buildup rates of the abstract and effector-selective decision signals on the random dot motion task, no such effects were observed on the contrast-change task. Thus, age-related drift rate reductions are likely due to task-specific deficits in sensory encoding affecting the quality of evidence entering the decision process. These findings are in keeping with the literature on the effects of age on sensory processing; while there is ample psychophysical and neurophysiological^54–56^ evidence to suggest that motion processing declines with increasing age, sensitivity to stimuli of relatively low spatial frequency and high contrast, such as the checkerboard stimuli used in the current study, appears to be preserved in older adults^57–59^. Thus, taken together, our neurophysiological data suggest that ageing does not lead to a fundamental decline in perceptual decision formation processes per se, but rather impacts on certain sensory inputs. An implication of these observations is that remediation strategies aimed at enhancing perceptual decisions in older adults should target those specific aspects of sensory processing that are subject to age-related decline.

One of the unexpected findings to emerge from the neural data was the early build-up of the CPP in older adults on the random dot motion task. We tested the hypothesis that this early build-up reflected greater premature (i.e. pre-target) accumulation in the older group by including starting point variability parameter as a free parameter in our model on the grounds that it would capture any variance in performance due to evidence accumulation during the inter-trial interval^42,60^. The addition of this parameter improved the fit to behaviour and also indicated higher levels of starting point variability in the older group, consistent with our CPP observations; because the CPP manifests as a positive deflection irrespective of the participant’s choice^61,62^, its amplitude at onset of coherent motion should be larger with higher levels of starting point variability. The reasons for this early buildup are unclear. One possibility is that older participants implemented a different evidence accumulation strategy in comparison to the younger group. Previous work from our lab has shown that in cases of temporal uncertainty of target onset, younger participants rely on early target selection signals to initiate neural evidence accumulation and these findings motivated our initial decision to commence the accumulation process in our models at the time of evidence onset. However, the premature buildup of the CPP exhibited by the older adults in the motion discrimination tasks suggests that it is unlikely that the decision process is started at evidence onset in this cohort. Rather, the current CPP data suggest that older adults accumulate evidence over the course of the inter-trial interval, perhaps owing to a reduced sensitivity to coherent motion onset. In this case, it is possible that both young and older participants implement a continuous “leaky” accumulation-to-bound process in which older samples of evidence are discounted as time passes^63^. The increased pre-target build-up of the CPP in the older adults could arise from reduced leakage, possibly as a strategic adaptation to avoid missed targets. While the introduction of starting point variability to our model allowed us to account for variations in behaviour and CPP arising from pretarget accumulation, further research will be required to establish the precise accumulation strategies that are implemented in continuous monitoring contexts.

Leveraging human brain signatures of decision formation to constrain computational signals has potential benefits that extend beyond research on ageing. Recent studies have suggested that the full drift diffusion model may be more complex than required in certain circumstances and this complexity can lead to more variable parameter estimates. The key message from these studies is that the sensitivity of diffusion models to between-group effects can be enhanced by reducing or constraining some of its parameters. However, determining which parameters should be constrained in a given experiment is not straightforward. For example, the EZ diffusion model^64^ does not include between-trial variability in any of its parameters and has been shown to be a more powerful tool at detecting simulated between-group effects as a result of this simplification^65^. Yet, our neurophysiological data suggest that its exclusion in the present context would overlook an important aspect of how ageing impacts the decision-making process. Here, we show that inspection of neurophysiological signals reflecting decision formation processes can provide a principled way of constraining the parameters of abstract decision models (see refs. ^25,28,36^ for similar approaches). Given the success of this approach in reconciling the differences in our behavioural and neural indices of decision making, this study paves the way for using similar techniques to address questions in clinically-relevant groups with impaired decision making, such as those with ADHD (e.g. ref. ^66^) and those suffering from addiction (e.g. ref. ^67^).

## Methods

### Participants

Thirty-nine younger and 42 older adults volunteered to take part in the study. The younger participants were recruited from the student population of Trinity College Dublin and compensated with research credits for their time, while older participants were recruited from the Trinity College Institute of Neuroscience volunteer participant panel. Criteria for inclusion in the study were right-handedness; normal or corrected-to-normal vision; no personal history of neurological or psychiatric illness, brain injury, abuse of substances or use of psychotropic drugs; no personal history or family history of epilepsy, unexplained fainting or sensitivity to flickering light; and a minimum score of 26 on the Mini Mental State Examination (MMSE). The age groups were matched for gender and years of education (Younger: 15.1, Older: 15.9), however, the younger group had a significantly higher average score on the MMSE than the older group (Young: 28.8, Older: 27.7, p<0.001). All participants were naive to the purposes of the study and provided written consent to participate. Participants were excluded if more than two-thirds of their trials were rejected due to EEG artifacts. This led to the rejection of two young participants and four older participants on the random dot motion task and one young participant and four older participants on the contrast-change detection task. A further two young and seven older participants were excluded from the random dot motion task due to technical issues during data acquisition. Following participant rejection for each experiment, there was a final sample of 35 younger adults (17 males, age range: 18-38, mean age: 22 years old) and 31 older adults (13 males, age range: 66-83, mean age: 74 years old) on the random dot motion task and a sample of 38 younger adults (15 males, age range: 18-38, mean age: 22 years old) and 38 older adults (16 males, age range: 66-85 years old, mean age: 74 years old) on the contrast-change detection task. All recruitment and experimental procedures were approved by the School of Psychology Research Ethics Committee, Trinity College Dublin in accordance with the principles of the Declaration of Helsinki.

### Procedure

Participants performed a continuous version of the random dot motion discrimination task^32^ and a contrast-change detection task^33^. Both tasks were completed in the same session in a darkened and sound-attenuated room, with the order of the tasks pseudorandomised across participants. In the same testing session, participants also participated in a visual oddball task, the results of which will be reported in a subsequent publication. Stimuli were presented on a 51 cm CRT monitor operating at 85 Hz and a resolution of 1024×768. Participants were seated at a distance of 55 cm from the display and were instructed to fixate a centrally-presented fixation point at all times during task performance. Prior to each task, participants carried out a practice block to familiarise themselves with the task and stimuli. During practice sessions, participants were given feedback on hits, misses and false alarms.

### Random dot motion task

Participants continuously monitored a patch of incoherently moving dots for intermittent targets defined by a period of coherent motion in the leftward or rightward direction (Figure 1a). Motion direction and coherence level (30% or 60%) were varied independently and randomly on a target-by-target basis. To facilitate the measurement of motor preparation signals, participants were asked to indicate leftward motion with a left-hand button press and rightward motion with a right-hand button press. Participants were instructed to avoid guessing and to respond as soon as they were certain they perceived coherent motion. The inter-target interval, during which the incoherent motion was continuously displayed, lasted 3, 5 or 7 seconds and was randomly chosen on a target-by-target basis.

Motion stimuli were presented within an 8-degree aperture centred on the fixation point and were displayed against a black background. During incoherent motion, an average of 150 white dots (each 6 × 6 pixels) were placed randomly and independently within the circular aperture on each of a sequence of 58.8 ms frames played at 17 frames per second. During coherent motion, a proportion of the dots were randomly selected on each frame to be displaced in either a leftward or rightward direction on the following frame at a speed of 6 degrees per second. In order to accommodate very slow responses from older participants, coherent motion targets were displayed for a maximum of 10 seconds or until 500 ms after the participant responded. Participants completed 6 blocks of the task, each lasting approximately 5 minutes and comprising 30 targets.

### Gradual contrast-change detection task

Participants continuously monitored a flickering (21.25Hz) annular checkerboard pattern for intermittent targets defined by a gradual decrease in contrast. The checkerboard stimulus (inner radius=3 degrees, outer radius=8 degrees) consisted of alternating light and dark radial segments presented against a dark grey background and was located in the centre of the display. Targets consisted of a linear decrease in contrast from 70 to 40% over a period of 1.6 seconds followed by a return to 70% contrast over a further 0.8 seconds (Figure 4s). The inter-target interval, during which the stimulus flickered at 70% contrast, lasted 3, 5 or 8 seconds and was randomly chosen on a target-by-target basis. Participants were instructed to avoid guessing and to make a mouse button press with their right index finger as soon as they were certain that the annulus was decreasing in contrast. Participants completed 4 blocks of the task, each lasting approximately 4 minutes and comprising 24 targets.

### EEG acquisition and preprocessing

Continuous EEG data were recorded using an ActiveTwo system (Biosemi, The Netherlands) from 64 scalp electrodes and digitised at 512 Hz. Electrodes were arranged using the standard 10/20 setup. Vertical and horizontal eye movements were recorded from four electro-occulogram (EOG) electrodes located above and below the left eye and at the left and right outer canthus, respectively. Data were analysed using custom-made scripts in MATLAB (MathWorks) drawing on EEGLAB routines for reading data files and for spherical interpolation of noisy channels^68^. EEG data were re-referenced offline to the average reference and low-pass filtered to 40Hz using a two-way least-squares FIR filter.

EEG data were segmented into epochs of −750 to 2500 ms and −750 to 2100 ms for the random dot motion and gradual contrast-change detection tasks, respectively. For the gradual contrast-change detection task, epochs were baseline-corrected relative to the average signal in the 250 ms interval preceding target onset. Inspection of the data from the random dot motion task revealed that, on average, the onset of the CPP of the older group occurred before the onset of coherent motion (see Supplementary Fig. 2), suggesting a tendency towards premature evidence accumulation onset due to temporal uncertainty of the stimulus onset. Epochs were therefore baseline-corrected to the average signal from −600 to −500 ms with respect to coherent motion onset. Trials were rejected if the bipolar vertical EOG signal (upper minus lower) exceeded an absolute value of 200 μV or if any scalp electrode signal exceeded 100 *μ*V within a window of 750 ms pre-stimulus to 150 ms post-response. EEG data were converted to current source density^69^ to increase spatial selectivity and to reduce the spatial blurring effect of volume conduction.

### Signal analysis

#### SSVEP

In the contrast-change detection task the contrast-dependent steady-state visual evoked potential (SSVEP) provided a cortical representation of the sensory evidence. The SSVEP was measured at a frequency of 21.25 Hz using the discrete Fourier transform over a window of exactly 10 cycles (470 ms) of the stimulus flicker frequency in order to attenuate spectral leakage. We first identified the region of maximum amplitude of the SSVEP on the grand-average scalp topography over 10 cycles of the SSVEP immediately prior to target onset. For each participant, the pretarget SSVEP amplitude was averaged across trials and normalised by dividing by the average amplitude at 21.25 Hz across all electrodes and trials. This normalisation step was required to account for inter-individual differences in the amplitude of the SSVEP ^70,71^. A further baseline correction step was then applied by dividing the normalised SSVEP at each electrode by the average amplitude measured across all frequencies. On the basis of the resulting topographies, the SSVEP was averaged over 6 electrodes centred on standard sites Oz and POz for both age groups.

The temporal evolution of the SSVEP was measured for each participant by calculating the average SSVEP amplitude across trials over a 10-cycle window at the start of the epoch and progressively moving the window forward by one sample (step size= 50 ms) until the SSVEP was calculated across the entire stimulus- and response-locked epochs. The SSVEP was then normalised relative to the 470 ms pretarget window for each participant.

#### CPP

The timecourse of evidence accumulation was indexed by a centro-parietal positivity (CPP) in the event-related potential^32,33^. Stimulus-locked CPP waveforms were generated for each participant by averaging the single-trial epochs defined in *EEG acquisition and preprocessing*. Response-locked CPPs were derived by extracting epochs from −900 to 300 ms relative to the time of the response, retaining the same prestimulus baseline intervals as the stimulus-locked waveforms. For both tasks and age groups, CPP amplitude and latency were measured at a single electrode centred on the region of maximum component amplitude identified in the grand-average response-locked scalp topography (standard site Pz). The peak magnitude of the response-locked CPP was calculated as the maximum voltage within the −100 ms to 100 ms window centred on the individual response time. The buildup rate of the response-locked CPP was measured as the slope of a straight line fitted to the unfiltered ERP waveform over a time window of −250 to −100 ms.

#### Mu/Beta

On the random dot motion task, motor preparation signals were indexed as a decrease in lateralised oscillatory activity in the Mu and Beta bands (8-30Hz) excluding the single frequency bin at precisely 17 Hz (stimulus flicker frequency). Oscillatory power in the Mu/Beta band was measured in the hemisphere contralateral to the hand used to indicate the direction of dot motion. Mu/Beta power was measured over electrodes C3 and CP3 for right responses and C4 and CP4 for left responses with the electrode sites selected on the basis of the grand-average response-locked scalp topographies. Contralateral signals for right and left responses were averaged to produce a single contralateral waveform for each motion coherence condition. The temporal evolution of the Mu/Beta power was measured via a sliding boxcar window of 412 ms with a 58 ms step size. For each participant, Mu/Beta amplitude was normalised relative to the average signal from −600 to −500 ms with respect to coherent motion onset.

On the contrast-change detection task, lateralised Mu/Beta power was measured as the oscillatory power in the 8-30 Hz range excluding the 21.25 Hz frequency bin (stimulus flicker frequency) over the standard left hemisphere motor site C3. The temporal evolution of the Mu/Beta power was measured via a sliding boxcar window of 470 ms with a 50 ms step size. For each participant, Mu/Beta amplitude was normalised relative to the 470 ms pretarget window. The slope of the response-locked Mu/Beta power was measured over a time window of −250 to 50 ms in the response-locked waveform, while the trough of the Mu/Beta power was measured within the - 100 ms to 100 ms window centred on the individual response time.

#### Alpha

The degree of attentional engagement during task performance was indexed by variability in pretarget alpha-band activity. For each participant, the average alpha amplitude (8-14 Hz) was calculated over the 707 ms (exactly 15 cycles of the SSVEP) interval prior to target onset across all 17 posterior electrodes. To assess the variability of alpha power between the two age groups, the coefficient of variation was calculated by dividing the standard deviation of pretarget alpha amplitude by the mean activity. The coefficient of variation is closely related to the standard deviation of a sample; however, it is not dependent on the sample mean^72^ and is therefore an appropriate measure for situations in which there is potentially a difference in sample means.

#### Bayes factor analysis

To determine the relative strength of evidence behind our approach of fixing parameters in the drift diffusion model based on our neural signals analysis, we conducted a Bayes factor analysis of the slope and amplitude measures of the CPP and Mu/Beta power for each task using JASP^73^. The Bayes factor overcomes some of the issues associated with null hypothesis significance testing by quantifying the relative likelihood of the data under the null versus the alternative hypothesis. Specifically, we calculated the Jeffrey, Zellner and Siow (JZS; see ref. ^74^) Bayes factor with an effect size of 1 to determine the strength of evidence in favour/against a group-level difference in the slope and peak amplitude of the neural signals of interest in each task. A JZS Bayes factor can be interpreted such that a value of three favours the null hypothesis three times more than the alternative hypothesis, while a value of one third favours the alternative three times more than the null.

### Drift diffusion modelling

Behavioural data from the gradual contrast-change detection and random dot motion tasks were fit with one-choice^43^ and two-choice^75^ drift diffusion models, respectively. Drift diffusion models assume that decisions are made through a noisy accumulation process in which sensory evidence is accumulated over time from a starting point, *z*, and a response is initiated when the cumulative evidence reaches the correct, *a*, or incorrect, *0*, response boundary (see Figure 2a). The rate of accumulation is called the drift rate, *v*, and is assumed to reflect the quality of information driving the decision process. The mean drift rate can vary across trials and this variability is assumed to be normally distributed with a standard deviation of *η*. There is also within-trial variability, or noise, in the evidence accumulation process. This allows that processes with the same mean drift rate terminate at different times, leading to a distribution of response times (RTs), and occasionally at the wrong boundary, leading to incorrect responses. The noise within a trial is also assumed to be normally distributed with a standard deviation of s and is fixed at 0.1 to scale the other parameters^76^. All non-decision related processing is accounted for by a single non-decision parameter, *ter*, that incorporates additive delays associated with sensory encoding and motor execution. For the contrast-change detection task, we assumed that the drift rate rose linearly over the duration of the target stimulus to reflect the ramping evidence signal (i.e. decreasing contrast) in this task. Guided by our neurophysiological observations from the motion discrimination task, we also introduced across-trial starting point variability (*sz*) into the model for that experiment and this was assumed to be uniformly distributed. Given the continuous nature of the experimental tasks, we made an additional assumption that if the evidence accumulation process had not terminated at one of the response boundaries by a time deadline, the trial was classified as a miss (see also ref. ^77^). The response deadlines were set to 10,000ms and 1750ms for the random dot motion and contrast-change detection tasks, respectively. We also made the additional assumption that the evidence accumulation process commenced following target onset. This decision was motivated by previous work from our lab showing that in scenarios involving temporal uncertainty of target onset, early target selection signals appear to play a role in initiating the neural evidence accumulation process^40^.

Unlike discrete versions of the random dot motion task, erroneous discriminations on the continuous version of the task were rare (~1% and 4% of targets in young and older participants, respectively). Several features of the data suggest that the few errors that did occur were likely not the result of evidence accumulation towards the incorrect decision boundary, but rather arose from erroneous action selection. First, older adults made significantly more errors when coherence was high than when it was low (5.4% vs. 2.6%, U=260, p<0.001, Mann-Whitney test), a pattern inconsistent with the predictions of the diffusion model (for example, see ref. ^75^). Second, error rates were greater than false alarms rates for both younger and older adults (Younger: 0.004 vs. 0.002 per second, p=0.01, Older: 0.015 vs. 0.004 per second, p<0.001, Wilcoxon signed rank test). Given that false alarms occur in conditions with no sensory evidence (i.e. drift rate=0) and errors occur with a positive drift rate diffusing towards the correct decision boundary, if errors reflected crossings of the incorrect decision boundary we would expect to see fewer errors than false alarms. Together these observations suggest that most of the errors we observe on the motion discrimination task do not arise from the evidence accumulation process itself. As a result, errors were excluded from the diffusion modelling analysis of the motion discrimination data.

We fit a number of diffusion models to the behavioural data from each perceptual task with varying parameter constraints and estimated parameter values by minimising the *G*^2^ statistic with a SIMPLEX minimisation routine. In order to fit the model to the data, five RT quantiles (0.1, 0.3, 0.5, 0.7, 0.9) were calculated from the RT distribution on correct trials and the proportion of trials lying between those quantiles were multiplied by the total number of trials to yield observed values. All RTs lying between 0 ms and the deadline in each task were included in these fits. These quantiles were then fed into the drift diffusion model to calculate the simulated proportion of trials that lay between these RT quantiles and were multiplied by the total number of trials to yield the model-derived expected values. The goodness-of-fit between the observed values and expected values was calculated via a *G*^2^ test and this statistic was minimised to provide estimates of the key model parameters. In models where parameters were constrained to reflect a lack of an age group difference in our neurophysiological observations, parameters were fixed to the mean of the young and older estimates from the unconstrained model fit to the pooled data. Model comparisons were performed using Bayes Information Criterion (BIC). The BIC provides a trade-off between model complexity and goodness-of-fit, favouring a model with less parameters if the differences in the degrees of freedom outweigh the gains associated with a better model fit. The preferred model for each task was chosen based on which produced the smallest BIC value.

For each experiment, we fit the model to the data in two ways. First, we pooled data across participants and fit the model separately to the younger and older group data. Here our approach was to find the most parsimonious version of the model (fewest number of parameters) to adequately fit the data in order to avoid overfitting (see ref. ^65^ for discussion of this topic). To this end, we first attempted to fit the pooled data with just the core components of the drift diffusion model free to vary (drift rate, decision boundary and non-decision time). However, this initial model did not provide a good fit to the data and through model simulations we identified that we also needed to include between-trial drift rate variability as a free parameter to capture the shape of the RT distributions. The addition of between-trial drift rate variability greatly reduced *G* values in both the younger and older data. Therefore, we fit this model to each participant’s data individually and averaged the resulting parameter values across participants. These recovered parameter values were then subject to inferential statistical analyses.

## Code availability

Custom code used for conducting EEG analysis and drift diffusion modelling available upon request to the corresponding author.

## Data availability

The data that support the findings of the study are available upon request to the corresponding author.

## Acknowledgements

This work was supported by a European Research Council (ERC) Starting Grant (to ROC) under the European Union’s Horizon 2020 research and innovation program (Grant Agreement No. 638289), by a Science Foundation Ireland ERC Support Grant (to ROC) and an Irish Research Council Postgraduate Scholarship (to AH). SPK is supported by a Career Development Award from Science Foundation Ireland (15/CDA/3591).

## Author contributions

ROC, SK and AH conceived and designed the experiments. AH collected the data. DPM analysed the data and fit the models. DPM, ROC and SK wrote the manuscript and all authors approved the final version.

## References

1. Levine, B., Svoboda, E., Hay, J. F., Winocur, G. & Moscovitch, M. Aging and autobiographical memory: dissociating episodic from semantic retrieval. Psychol. Aging 17, 677–689 (2002).

2. Gazzaley, A., Cooney, J. W., Rissman, J. & D’Esposito, M. Top-down suppression deficit underlies working memory impairment in normal aging. Nat. Neurosci. 8, 1298–1300 (2005).

3. Salthouse, T. A. Constraints on theories of cognitive aging. Psychon. Bull. Rev. 3, 287–299 (1996).

4. Wasylyshyn, C., Verhaeghen, P. & Sliwinski, M. J. Aging and task switching: a meta-analysis. Psychol. Aging 26, 15–20 (2011).

5. Stern, Y. What is cognitive reserve? Theory and research application of the reserve concept. J. Int. Neuropsychol. Soc. 8, 448–460 (2002).

6. Park, D. C. & Reuter-Lorenz, P. The adaptive brain: aging and neurocognitive scaffolding. Annu. Rev. Psychol. 60, 173–196 (2009).

7. Laming, D. Information theory of choice-reaction times. (London: Academic Press, 1968).

8. Link, S. W. & Heath, R. A. A sequential theory of psychological discrimination. Psychometrika 40, 77–105 (1975).

9. Ratcliff, R. A Theory of Memory Retrieval. Psychol. Rev. 85, 59–108 (1978).

10. Ratcliff, R., Thapar, A. & McKoon, G. The effects of aging on reaction time in a signal detection task. Psychol. Aging 16, 323–341 (2001).

11. Ratcliff, R., Thapar, A. & McKoon, G. A diffusion model analysis of the effects of aging on brightness discrimination. Percept. Psychophys. 65, 523–535 (2003).

12. Ratcliff, R., Thapar, A. & McKoon, G. Aging, practice, and perceptual tasks: a diffusion model analysis. Psychol. Aging 21, 353–371 (2006).

13. Starns, J. J. & Ratcliff, R. The effects of aging on the speed-accuracy compromise: Boundary optimality in the diffusion model. Psychol. Aging 25, 377–390 (2010).

14. Ratcliff, R., Thapar, A. & McKoon, G. Individual differences, aging, and IQ in two-choice tasks. Cognit. Psychol. 60, 127–157 (2010).

15. Spaniol, J., Voss, A. & Grady, C. L. Aging and emotional memory: cognitive mechanisms underlying the positivity effect. Psychol. Aging 23, 859–872 (2008).

16. Forstmann, B. U. et al. The speed-accuracy tradeoff in the elderly brain: a structural model-based approach. J. Neurosci. 31, 17242–17249 (2011).

17. Rabbitt, P. How Old and Young Subjects Monitor and Control Responses for Accuracy and Speed. Br. J. Psychol. 70, 305–11 (1979).

18. Ratcliff, R., Thapar, A., Gomez, P. & McKoon, G. A diffusion model analysis of the effects of aging in the lexical-decision task. Psychol. Aging 19, 278–289 (2004).

19. Ratcliff, R., Thapar, A. & McKoon, G. Aging and individual differences in rapid two-choice decisions. Psychon. Bull. Rev. 13, 626–635 (2006).

20. Ratcliff, R., Thapar, A. & McKoon, G. Application of the diffusion model to two-choice tasks for adults 75-90 years old. Psychol. Aging 22, 56–66 (2007).

21. Thapar, A., Ratcliff, R. & McKoon, G. A diffusion model analysis of the effects of aging on letter discrimination. Psychol. Aging 18, 415–429 (2003).

22. Ratcliff, R., Thapar, A. & McKoon, G. Effects of aging and IQ on item and associative memory. J. Exp. Psychol. Gen. 140, 464–487 (2011).

23. Dully, J., McGovern, D. P. & O’Connell, R. G. The impact of natural aging on computational and neural indices of perceptual decision making: A review. Behav. Brain Res. (2018). doi:10.1016/j.bbr.2018.02.001

24. Robertson, I. H. A noradrenergic theory of cognitive reserve: implications for Alzheimer’s disease. Neurobiol. Aging 34, 298–308 (2013).

25. Hanks, T., Kiani, R. & Shadlen, M. N. A neural mechanism of speed-accuracy tradeoff in macaque area LIP. eLife 3, (2014).

26. Hanks, T. D., Mazurek, M. E., Kiani, R., Hopp, E. & Shadlen, M. N. Elapsed decision time affects the weighting of prior probability in a perceptual decision task. J. Neurosci. 31, 6339–6352 (2011).

27. Heitz, R. P. & Schall, J. D. Neural chronometry and coherency across speed-accuracy demands reveal lack of homomorphism between computational and neural mechanisms of evidence accumulation. Philos. Trans. R. Soc. Lond. B. Biol. Sci. 368, 20130071 (2013).

28. Purcell, B. A. & Kiani, R. Neural Mechanisms of Post-error Adjustments of Decision Policy in Parietal Cortex. Neuron 89, 658–671 (2016).

29. Hanes, D. P. & Schall, J. D. Neural control of voluntary movement initiation. Science 274, 427–430 (1996).

30. Roitman, J. D. & Shadlen, M. N. Response of neurons in the lateral intraparietal area during a combined visual discrimination reaction time task. J. Neurosci. 22, 9475–9489 (2002).

31. Ratcliff, R., Hasegawa, Y. T., Hasegawa, R. P., Smith, P. L. & Segraves, M. A. Dual diffusion model for single-cell recording data from the superior colliculus in a brightness-discrimination task. J. Neurophysiol. 97, 1756–1774 (2007).

32. Kelly, S. P. & O’Connell, R. G. Internal and External Influences on the Rate of Sensory Evidence Accumulation in the Human Brain. J. Neurosci. 33, 19434–19441 (2013).

33. O’Connell, R. G., Dockree, P. M. & Kelly, S. P. A supramodal accumulation-to-bound signal that determines perceptual decisions in humans. Nat. Neurosci. 15, 1729–1735 (2012).

34. de Lange, F. P., Rahnev, D. A., Donner, T. H. & Lau, H. Prestimulus oscillatory activity over motor cortex reflects perceptual expectations. J. Neurosci. 33, 1400–1410 (2013).

35. Donner, T. H., Siegel, M., Fries, P. & Engel, A. K. Buildup of choice-predictive activity in human motor cortex during perceptual decision making. Curr. Biol. 19, 1581–1585 (2009).

36. Murphy, P. R., Boonstra, E. & Nieuwenhuis, S. Global gain modulation generates time-dependent urgency during perceptual choice in humans. Nat. Commun. 7, 13526 (2016).

37. Twomey, D. M., Kelly, S. P. & O’Connell, R. G. Abstract and Effector-Selective Decision Signals Exhibit Qualitatively Distinct Dynamics before Delayed Perceptual Reports. J. Neurosci. 36, 7346–7352 (2016).

38. Ball, K. & Sekuler, R. Improving visual perception in older observers. J. Gerontol. 41, 176–182 (1986).

39. Billino, J., Bremmer, F. & Gegenfurtner, K. R. Differential aging of motion processing mechanisms: evidence against general perceptual decline. Vision Res. 48, 1254–1261 (2008).

40. Loughnane, G. M. et al. Target Selection Signals Influence Perceptual Decisions by Modulating the Onset and Rate of Evidence Accumulation. Curr. Biol. 26, 496–502 (2016).

41. Steinemann, N. A., O’Connell, R. G. & Kelly, S. P. Decisions are expedited through multiple neural adjustments spanning the sensorimotor hierarchy. Nat. Commun. (In press).

42. Jepma, M., Wagenmakers, E.-J. & Nieuwenhuis, S. Temporal expectation and information processing: a model-based analysis. Cognition 122, 426–441 (2012).

43. Ratcliff, R. & Van Dongen, H. P. A. Diffusion model for one-choice reaction-time tasks and the cognitive effects of sleep deprivation. Proc. Natl. Acad. Sci. U. S. A. 108, 11285–11290 (2011).

44. Di Russo, F. et al. Spatiotemporal analysis of the cortical sources of the steady-state visual evoked potential. Hum. Brain Mapp. 28, 323–334 (2007).

45. Hanslmayr, S. et al. Prestimulus oscillations predict visual perception performance between and within subjects. NeuroImage 37, 1465–1473 (2007).

46. O’Connell, R. G. et al. Uncovering the neural signature of lapsing attention: electrophysiological signals predict errors up to 20 s before they occur. J. Neurosci. 29, 8604–8611 (2009).

47. Dockree, P. M. et al. The Effects of Methylphenidate on the Neural Signatures of Sustained Attention. Biol. Psychiatry 82, 687–694 (2017).

48. Murphy, P. R., Vandekerckhove, J. & Nieuwenhuis, S. Pupil-Linked Arousal Determines Variability in Perceptual Decision Making. PLOS Comput. Biol. 10, e1003854 (2014).

49. Turner, B. M., van Maanen, L. & Forstmann, B. U. Informing cognitive abstractions through neuroimaging: The neural drift diffusion model. Psychol. Rev. 122, 312–336 (2015).

50. Turner, B. M., Forstmann, B. U., Love, B. C., Palmeri, T. J. & Van Maanen, L. Approaches to analysis in model-based cognitive neuroscience. J. Math. Psychol. 76, 65–79 (2017).

51. Turner, B. M. et al. A Bayesian framework for simultaneously modeling neural and behavioral data. NeuroImage 72, 193–206 (2013).

52. Turner, B. M., Rodriguez, C. A., Norcia, T. M., McClure, S. M. & Steyvers, M. Why more is better: Simultaneous modeling of EEG, fMRI, and behavioral data. NeuroImage 128, 96–115 (2016).

53. Frank, M.J. et al. fMRI and EEG Predictors of Dynamic Decision Parameters during Human Reinforcement Learning. J. Neurosci. 35, 485 (2015).

54. Yang, Y. et al. Aging affects contrast response functions and adaptation of middle temporal visual area neurons in rhesus monkeys. Neuroscience 156, 748–757 (2008).

55. Yang, Y. et al. Aging affects the neural representation of speed in Macaque area MT. Cereb. Cortex 19, 1957–1967 (2009).

56. Liang, Z. et al. Aging affects the direction selectivity of MT cells in rhesus monkeys. Neurobiol. Aging 31, 863–873 (2010).

57. Owsley, C., Sekuler, R. & Siemsen, D. Contrast sensitivity throughout adulthood. Vision Res. 23, 689–699 (1983).

58. Elliott, D., Whitaker, D. & MacVeigh, D. Neural contribution to spatiotemporal contrast sensitivity decline in healthy ageing eyes. Vision Res. 30, 541–547 (1990).

59. Habak, C. & Faubert, J. Larger effect of aging on the perception of higher-order stimuli. Vision Res. 40, 943–950 (2000).

60. Laming, D. Choice reaction performance following an error. Acta Psychol. (Amst.) 43, 199–224 (1979).

61. Kelly, S. P. & O’Connell, R. G. The neural processes underlying perceptual decision making in humans: Recent progress and future directions. J. Physiol.-Paris 109, 27–37 (2015).

62. Afacan-Seref, K., Steinemann, N. A., Blangero, A. & Kelly, S. P. Dynamic Interplay of Value and Sensory Information in High-Speed Decision Making. Curr. Biol. 28, 795–802.e6 (2018).

63. Usher, M. & McClelland, J. L. The time course of perceptual choice: the leaky, competing accumulator model. Psychol. Rev. 108, 550–592 (2001).

64. Wagenmakers, E.-J., van der Maas, H. L. J. & Grasman, R. P. P. P. An EZ-diffusion model for response time and accuracy. Psychon. Bull. Rev. 14, 3–22 (2007).

65. van Ravenzwaaij, D., Donkin, C. & Vandekerckhove, J. The EZ diffusion model provides a powerful test of simple empirical effects. Psychon. Bull. Rev. 24, 547–556 (2017).

66. Hauser, T. U., Fiore, V. G., Moutoussis, M. & Dolan, R. J. Computational Psychiatry of ADHD: Neural Gain Impairments across Marrian Levels of Analysis. Trends Neurosci. 39, 63–73 (2016).

67. Bechara, A. Decision making, impulse control and loss of willpower to resist drugs: a neurocognitive perspective. Nat. Neurosci. 8, 1458–1463 (2005).

68. Delorme, A. & Makeig, S. EEGLAB: an open source toolbox for analysis of single-trial EEG dynamics including independent component analysis. J. Neurosci. Methods 134, 9–21 (2004).

69. Kayser, J. & Tenke, C. E. Principal components analysis of Laplacian waveforms as a generic method for identifying ERP generator patterns: I. Evaluation with auditory oddball tasks. Clin. Neurophysiol. 117, 348–368 (2006).

70. Silberstein, R. B. et al. Steady-state visually evoked potential topography associated with a visual vigilance task. Brain Topogr. 3, 337–347 (1990).

71. Silberstein, R. B., Nunez, P. L., Pipingas, A., Harris, P. & Danieli, F. Steady state visually evoked potential (SSVEP) topography in a graded working memory task. Int. J. Psychophysiol. 42, 219–232 (2001).

72. Barlow, J. S. The Electroencephalogram: Its Patterns and Origins. (MIT Press, 1993).

73. JASP team. JASP (Version 0.8.3) [Computer software]. (2017).

74. Rouder, J. N., Speckman, P. L., Sun, D., Morey, R. D. & Iverson, G. Bayesian t tests for accepting and rejecting the null hypothesis. Psychon. Bull. Rev. 16, 225–237 (2009).

75. Ratcliff, R. & McKoon, G. The diffusion decision model: theory and data for two-choice decision tasks. Neural Comput. 20, 873–922 (2008).

76. Ratcliff, R. & Tuerlinckx, F. Estimating parameters of the diffusion model: approaches to dealing with contaminant reaction times and parameter variability. Psychon. Bull. Rev. 9, 438–481 (2002).

77. Murphy, P. R., Robertson, I. H., Harty, S. & O’Connell, R. G. Neural evidence accumulation persists after choice to inform metacognitive judgments. eLife 4, (2015).

